# A Taxonomy of Seizure Spread Patterns, Speed of Spread, and Associations With Structural Connectivity

**DOI:** 10.1101/2022.10.24.513577

**Authors:** Andrew Y. Revell, Akash R. Pattnaik, Erin Conrad, Nishant Sinha, Brittany H. Scheid, Alfredo Lucas, John M. Bernabei, John Beckerle, Joel M. Stein, Sandhitsu R. Das, Brian Litt, Kathryn A. Davis

## Abstract

Although seizure detection algorithms are widely used to localize seizure onset on intracranial EEG in epilepsy patients, relatively few studies focus on seizure activity beyond the seizure onset zone to direct treatment of surgical patients with epilepsy. To address this gap, we develop and compare fully automated deep learning algorithms to detect seizure activity on single channels, effectively quantifying spread when deployed across multiple channels. Across 275 seizures in 71 patients, we discover that the extent of seizure spread across the brain and the timing of seizure spread between temporal lobe regions is associated with both surgical outcomes and the brain’s structural connectivity between temporal lobes. Finally, we uncover a hierarchical structure of seizure spread patterns highlighting the relationship between clusters of seizures. Collectively, these findings underscore the broad utility in quantifying seizure activity past seizure onset to identify novel mechanisms of seizure evolution and its relationship to potential seizure freedom.

## Introduction

Seizure onset, timing, extent of activity, and other patterns of seizure activity captured during a seizure are used in the clinical interpretation of EEG to plan treatment for refractory epilepsy ^1–5^. Surgical removal of epileptogenic tissue through resection or ablation may be appropriate given sufficient clinical evidence that the removal of localized brain tissue can cure a patient of epilepsy or improve their quality of life ^6,7^. In other cases, patterns captured on EEG may instead indicate other treatment modalities, such as implantable neuromodulatory devices ^8,9^, or other palliative options ^10,11^.

Correct identification of the seizure onset zone and its surgical removal offers the best chance for complete seizure freedom for patients with refractory epilepsy ^12–17^. Accurate localization of the seizure onset has thus been a primary focus in epilepsy research to improve outcomes. Yet, overall seizure freedom rates after surgery have remained relatively stagnant over the last 30-40 years and vary greatly across centers and studies quantifying outcomes ^6^.

To improve outcomes of refractory epilepsy patients, the focus in epilepsy research perhaps should also include efforts in identifying patterns of seizure activity beyond seizure onset — a seizure’s timing, speed, extent of activity, and spread may be just as important in identifying distinct pathophysiological mechanisms of seizure evolution and the best course of treatment for a patient with a specific type (or types) of seizure spread patterns. However, we currently lack fully automated and validated measures to quantify the spread of seizure activity.

Here, we develop and compare deep learning algorithms with simple features to quantify seizure spread in 71 patients across 275 seizures. We use the best performing algorithm to answer three main questions: (1) Is the extent and timing of spread associated with patient outcomes? (2) Is the timing of seizure spread related to the structural connectivity of the brain? (3) What are the rules governing seizure spread — is there a hierarchical organization separating the patterns of seizure spread into distinct clusters while grouping related seizures across patients together?

## Results

### A. Deep Learning Algorithms are Effective in Differentiating Ictal and Interictal States

To investigate the hierarchical organization of seizure spread patterns across seizures and patients, we need robust measures of seizure spread. Currently, a limited number of studies deploy automated algorithms to quantify seizure spread and usually rely on single features, such as line length ^18^ or power ^2^; however, we did not know if such algorithms reliably measure seizure spread. Many studies that do quantify spread are performed with a small number of patients or require manual annotations by an epileptologist ^19,20^. We compare the performance of different large-scale, and completely automated seizure detection algorithms to capture spread. We use both simple EEG features (absolute slope, line length, and broadband power) and three deep learning algorithms with different neural network architectures designed for time-series data.

We chose the single EEG features because they have been shown to correspond with clinical annotations for seizure onset ^17,21–23^ and they are a relatively small number of simple features to compare and preserve power in our study. The deep learning algorithms were chosen because they are effective predictors of time series data ^24^. The deep learning algorithms were (1) WaveNet ^25^, a one-dimensional conventional neural network (1D CNN) with a causal and dilated neural network architecture, (2) a 1D CNN, denoted here as a default CNN as opposed to WaveNet with a specific neural network architecture, and (3) a long-short-term-memory (LSTM) neural network originally applied for sequence modeling ^26^.

To measure spread, we first trained the deep learning algorithms to differentiate between two states, ictal and interictal states, so that we could eventually quantify *when* the state transition happens across channels (Fig 2a). The area under the curve (AUC) was calculated for differentiating the two states with a leave-one-patient-out (n = 13 patients) cross validation across varying learning rates (Fig 2b). Similarly, a leave-one-out cross validation was performed on the single features for differentiating ictal and interictal states (Fig 2c). The deep learning algorithms at a default learning rate of 0.001 outperform the AUC of the single features (p < 0.001, Wilcoxon Signed-rank test, two sided, FDR correction for 15 tests pairwise across the 6 algorithms), except the comparison between LSTM and power (p > 0.05).

**Fig. 1.**
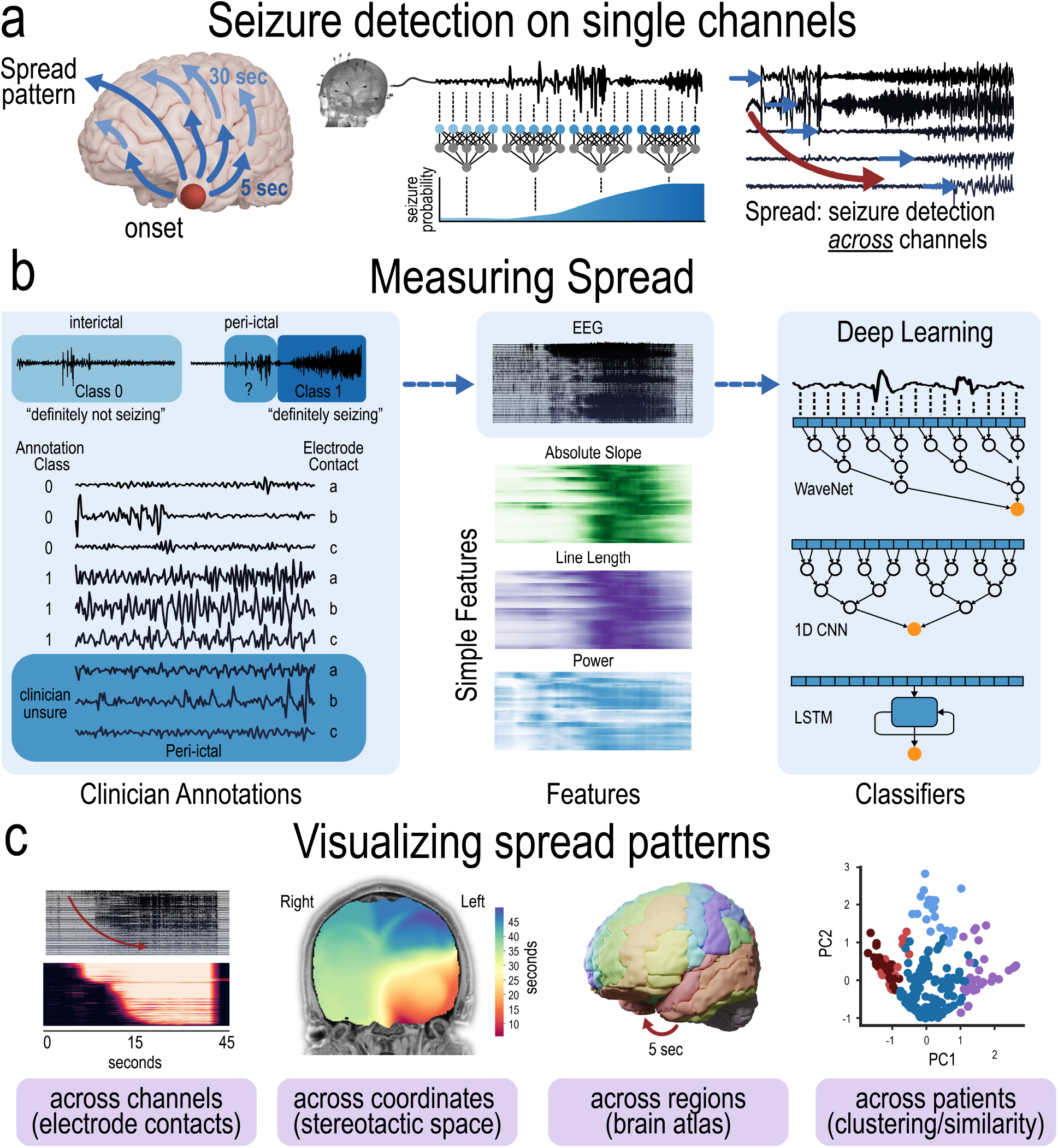
Seizure detection on single channels. **a**, Seizure spread is quantified by measuring seizure activity across multiple channels. A pattern of spread is characterized by the extent, timing, speed, and locations of seizure activity. **b**, Schematic showing how seizure spread can be measured. Machine learning algorithms can be deployed to differentiate two classes: definitely seizing (ictal) states, and definitely not seizing (interictal) states. Once excellent performance has been achieved to differentiate states, fully automated algorithms can be deployed to determine state transitions on peri-ictal data. Simple features such as absolute slope, line length, and power — all associated with seizure onset — can be used. EEG data can also be used in the case of deep learning algorithms such as (1) WaveNet, a one-dimensional conventional neural network (1D CNN) with a causal and dialated neural network architecture, (2) a 1D CNN, denoted here as a default CNN as opposed to WaveNet with a tailored architecture, or (3) a long-short-term-memory (LSTM) neural network. **c**, Seizure spread can be visualized in four different ways, all aided to enhance our understanding of seizure spread patterns.

**Fig. 2.**
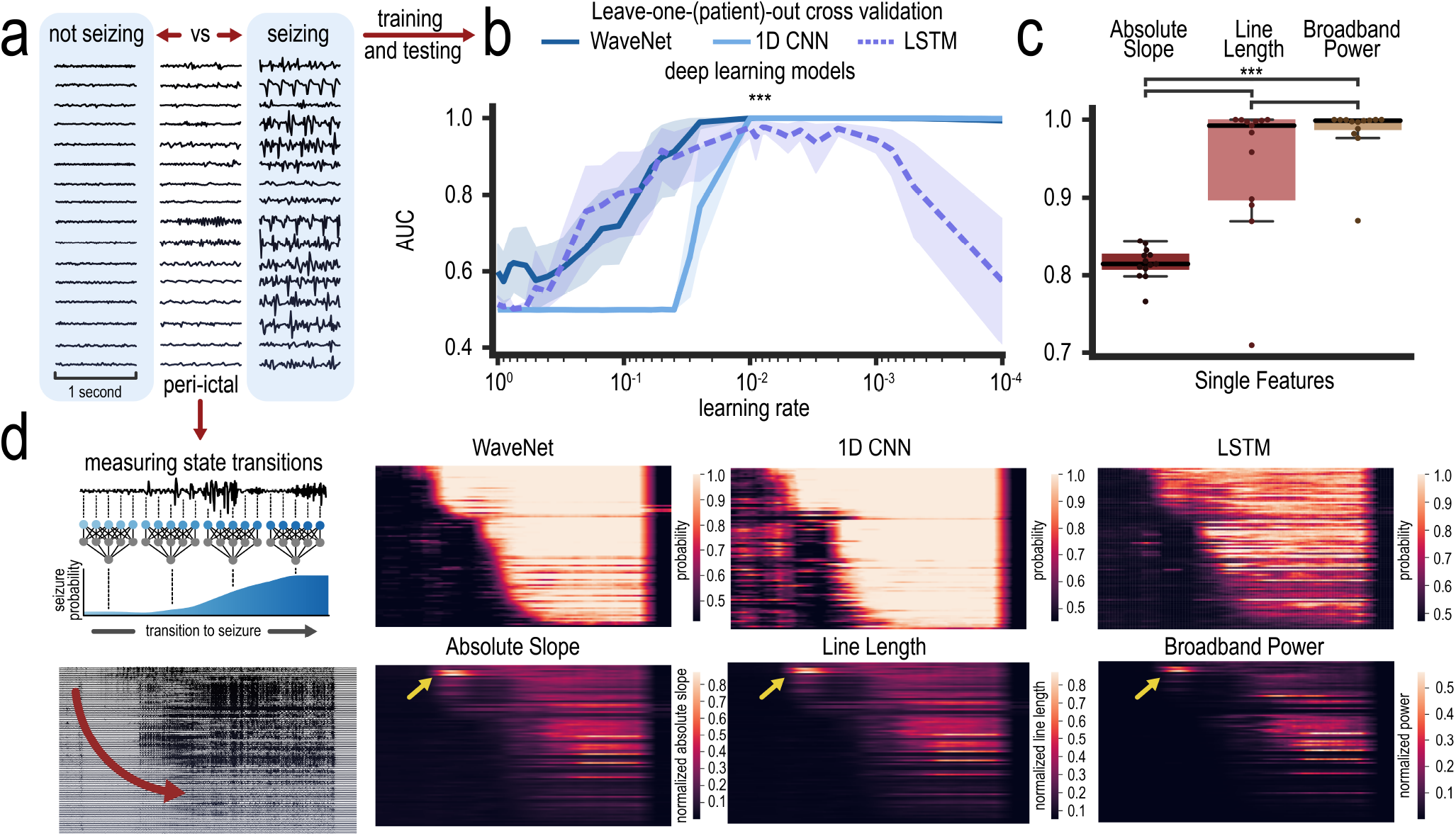
Training and Testing of Binary Seizure States to Measure State Transitions. **a**, Examples of one-second windows of non seizing (interictal > 6 hours before seizure) and seizing states of the same channel in each row at the same gain. These two states were used for training and testing of deep learning algorithms to be deployed on peri-ictal data nearing the transition to a seizure state. Peri-ictal data show windows 1-20 seconds before seizure states of the same channel. **b**, Leave-one-out (n = 13 patients) cross validation and AUC as a function of learning rate is shown for the three deep learning algorithms. At a default learning rate of 0.001, the AUC was compared with each of the single feature AUC (*** p < 0.001, Wilcoxon Signed-rank test, two sided, FDR correction for 15 tests pairwise across the 6 algorithms). Shaded areas represent 95% CIs. **c**, Leave-one-out (n = 13 patients) cross validation and AUC for each of the three single feature algorithms in detecting seizing vs non seizing binary states. (*** p < 0.001 (Wilcoxon Signed-rank test, two sided, FDR correction for 15 tests pairwise across the 6 algorithms). **d**, Schematic showing that these algorithms were deployed on Peri-ictal data to measure state transitions. Heatmap and colorers indicate seizure probabilities (for the deep learning algorithm) or normalized feature values (for the single features) to measure seizure spread across time (x-axis) and across channels (y-axis, order is the same across heat maps) of the example seizure shown at the bottom left. Yellow arrows point to seizure onset channels the single feature pick up, however, the pattern of activity as shown in the heatmap is not similar between the algorithms.

Once the algorithms are developed to differentiate ictal and interictal states for each channel, they can be deployed on peri-ictal data to measure the time that the state transition occurs. The timing of state transitions across multiple channels effectively measures the spread of seizure activity across the brain (Fig 2d). In the case of the deep learning algorithms, the transition from interictal to ictal states occurs when the probability of an ictal state surpasses a set threshold. In the case of single features, the transition from interictal to ictal states occurs when each respective normalized feature (rather than probabilities) surpasses a set threshold. The deep learning algorithms have larger contrasts between inter-ictal and ictal states than the single features (Fig 2d).

### B. Deep Learning Algorithms Outperform Single Features in Detecting Seizure Onset Contacts

To validate the algorithms measuring seizure spread, we first examine their performances on detecting the initial spread points – the seizure onset contacts (Fig 3a).

**Fig. 3.**
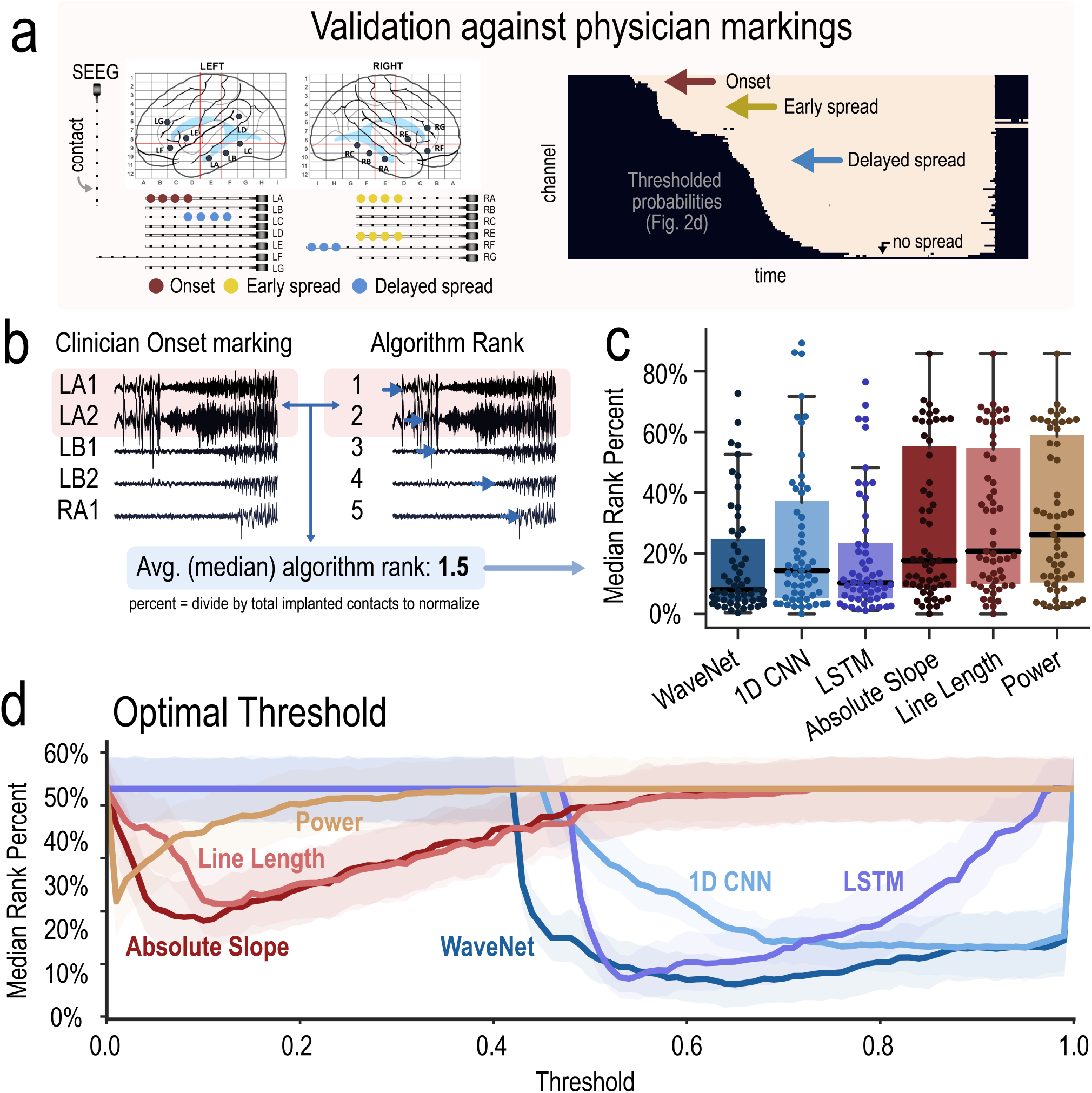
Validation of Seizure Spread Detection Algorithms. **a**, A patient example with physician markings is shown with a corresponding thresholded activation map from the WaveNet algorithm. **b**, This schematic shows how the agreement between physician markings of seizure onset contacts (red) and each algorithm marking is calculated. The algorithm ranking of the seizure onset contacts were averaged (median) and divided by the total number of implanted contacts to normalize for differences in implantations. Note, this calculation penalizes ranking scores for physicians who marked large number of contacts. **c**, Box plots showing the median rank percent for all patients with seizure onset contact annotations (n = 55). All six algorithms perform better than chance (one-sample Wilcoxon signed rank test, two sided, FDR correction for 6 tests, null hypothesis is 50% median rank – if a seizure onset contact is randomly assigned a rank, it would be 50% of all implanted contacts). WaveNet, 1D CNN, LSTM all perform better than the single feature algorithms (P < 0.01, Wilcoxon signed rank test, two sided, FDR correction for 15 tests pairwise across the 6 algorithms), but none of the deep learning algorithms outperform each other. **d**, The optimal threshold for each algorithm in agreement with physician markings of seizure onset. The optimal thresholds were chosen for the analysis in panel c.

The agreement between physician markings and each algorithm marking is calculated. For example, a clinician may mark LA1 and LA2 as the seizure onset contacts (Fig 3b). The seizure spread algorithm also independently determines the rank order of seizure activity for all contacts. The rank order from the algorithm is averaged (by computing the median) for just the clinician onset contacts. In other words, if the algorithm determines that contact LA1 started to seize first and contact LA2 started to seize second, the median rank of these contacts is 1.5. This median ranking is divided by the total number of implanted contacts to normalize for differences in number of contacts across patients.

All six algorithms perform better than chance in detecting seizure onset contacts (Fig 3c), p < 0.05, one-sample Wilcoxon signed rank test, two sided, FDR correction for 6 tests, null hypothesis is 50% median rank – if a seizure onset contact is randomly assigned a rank, it would be 50% of all implanted contacts).

All three deep learning algorithms perform better than the single feature algorithms in differentiating states (p < 0.01, Wilcoxon signed rank test, two sided, FDR correction for 15 tests pairwise across the 6 algorithms), but none of the deep learning algorithms outperform each other.

We also calculated the performance of each algorithm at varying thresholds (Fig 3d). The optimal threshold for each deep learning algorithm is a probability of 0.69 (WaveNet), 0.60 (1D CNN), 0.94 (LSTM), 0.26 (absolute slope), 0.11 (line length), and 0.02 (broadband power). The comparison in Fig 3c was made at each algorithm’s optimal threshold – the deep learning algorithms capture relevant seizure onset contacts across more patients and across wider ranges of thresholds than the single features (i.e. the deep learning algorithms may more likely capture relevant seizure spread patterns without having to tune specific threshold parameters).

### C. The Extent of Seizure Spread – Poor Outcome Patients Have More Distributed Regions Involved in Seizures

The extent of seizure spread over its evolution can be quantified in two ways: (i) by the number of contacts activated over time and (ii) by the number of brain regions activated over time (Fig 4).

**Fig. 4.**
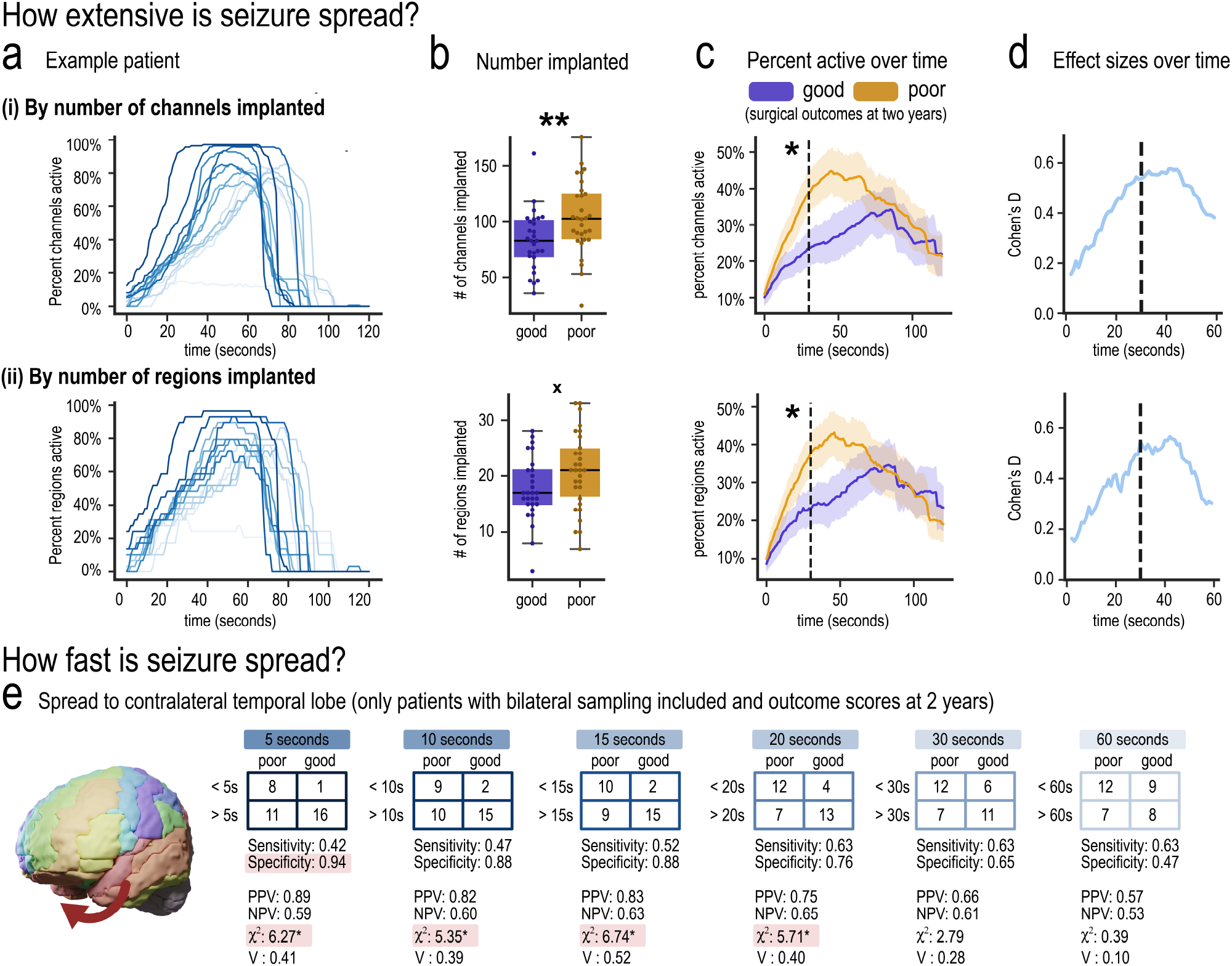
Extent and Speed of Seizure Spread. The extent of seizure spread over time is quantified by (i) the number of channels and (ii) the number of regions active. **a**The percentage of channels and regions active over time for a patient with 14 seizures is shown. Darker lines indicate seizures captured earlier during their hospital stay. **b**, Extent of seizure spread can be biased by the number of contacts or regions implanted. Poor outcome patients (n = 30) have higher number of contacts implanted than good outcome patients (n = 28, ** p < 0.01 Mann Whitney U test, two sided, null hypothesis: number of contacts implanted is equal between outcomes). However, poor outcome patients (n = 30) did not have significantly higher number of regions sampled than good outcome patients (n = 28, x denotes trending p < 0.10, Mann Whitney U test, two-sided, null hypothesis: number of sampled regions is equal between outcomes). **c**, Even with larger sampling (more contacts), poor outcome patients (n = 30) have higher percentage of contacts active than good outcome patients (n = 28) at the 30 second mark (dashed line, * p < 0.05, Mann Whitney U test, two-sided, null hypotheses: percentage of active contacts at 30 seconds is equal between outcomes). Similarly, poor outcome patients have higher percentages of sampled regions active than good outcome patients (n = 28) at 30 seconds (p < 0.05, Mann Whitney U test, two-sided, null hypothesis: percentage of active implanted regions at 30 seconds is equal between good and poor outcome patients). Shaded areas represents 68% CIs. **d**, Effect sizes of the differences between outcomes is shown over time. Dashed line is at 30 seconds. **e**, Contingency tables comparing the speed of spread between temporal lobes in good and poor outcomes. Sensitivity, specificity, positive predictive values (PPV), negative predictive values (NPV), chi-square test, and Cramer’s V are reported for each cutoff. * p < 0.05, FDR correction for 6 tests for the 6 cutoffs (null hypothesis: no association between timing of spread at a specific cutoff and outcome. Red highlights indicate cutoffs with significant associations or high specificity.

The percent of contacts and regions activated over time in an example patient with 14 seizures is shown in (Fig 4a) using the WaveNet algorithm at its optimal threshold. Other algorithms are shown in (Fig S2). Darker lines represent seizures captured earlier during their hospital stay. Earlier seizures have more rapid activation (larger slopes) of contacts and regions. The velocity of activation has a noticeable shift with smaller slopes and longer seizures at approximately the 6-8th seizure. Here, we see evidence that the the evolution of seizures across time and across seizures themselves can change – the quality of the seizures within a patient can change during their hospital course and may be due to a variety of factors (for example, medication changes). In other words, the changes in seizure patterns within a patient can be captured with algorithms designed to quantify seizure spread (are their seizures stereotypical? Is their pattern changing? Should we use this seizure to localize seizure onset for surgery given its stereotypical nature?).

We also hypothesized that the pattern of seizure spread between good and poor outcome patients is different. We reasoned that poor outcome patients may have more distributed (extensive) regions involved during the seizure ^27^. Before testing this hypothesis, however, we reasoned the extent of seizure spread can be biased by a different number of contacts and regions targeted for implantation between the two groups (Fig 4b). We found that poor outcome patients (n = 30) typically had higher number of *contacts* implanted than good outcome patients (n = 28, * p < 0.01, Mann Whiteney U test, null hypothesis: the number of implanted contacts is the same between good and poor outcome patients). In contrast, we found that poor outcome patients (n = 30) did not have significantly higher number of brain *regions* sampled than good outcome patients (n = 28, x indicates trending p < 0.10, Mann Whiteney U test, null hypotheses: the number of sampled regions is the same between good and poor outcome patients).

Despite having more contacts and a similar number of sampled regions, poor outcome patients demonstrated a higher *percentage* of contacts and regions active over time (Fig 4c). In other words, poor outcome patients have more distributed (extensive) regions involved during their seizures despite a bias that would be expected to decrease the percentage of contacts or regions activated over time (poor outcome patients have more contacts implanted, and if the extent of seizure spread was equal between good and poor outcome patients, then the *percentage* of contacts activated would be less).

We also calculated effect size differences across time between good and poor outcome patients (Fig 4d). Effect sizes are largest between 20 – 50 seconds into a seizure indicating that the prediction of outcome using the extent of seizure activity may be best in this time window. However, effect sizes are low-to-moderate (< 0.8) and the percentages of active contacts or regions may not be sufficient metrics for clinical use. These results provide evidence that the pattern of spread may be different between good and poor outcome epilepsy patients. The observed pattern of spread may help direct treatment and indicate if surgical intervention may result in seizure freedom at two years.

### D. The Speed of Seizure Spread – Poor Outcome Patients Have Quicker Spread Between Temporal Lobe Regions

Fig 4e shows contingency tables comparing the speed of spread between temporal lobes in good and poor outcome surgical epilepsy patients. The activation time of all contacts in the temporal lobe structures are averaged together for each of the left and right hemispheres. The difference between these average activation times between the left and right lobes are recorded. Only patients with bilateral temporal lobe sampling and patients with outcome scores at 2 years are used. Cutoff spread times of 5, 10, 15, 20, 30, and 60 seconds are used to differentiate good and poor outcome patients. Sensitivity, specificity, positive predictive values (PPV), negative predictive values (NPV), chi-square test, and Cramer’s V are reported for each cutoff.

We show that there is an association between timing of spread and surgical outcome if seizures spread between temporal lobes within 5, 10, 15, and 20 seconds (p < 0.05, FDR correction for 6 tests for the 6 cutoffs, chi-squared test, null hypothesis: there is no association between the timing of spread at a specific cutoff and surgical outcome). Spread within 5 seconds has the highest specificity (94%), thus patients with seizures that spread quickly have a high likelihood of a poor outcome. Speed of spread at any time does not provide good sensitivity (40-60%, i.e. many patients with slow or no spread still have poor outcomes).

Cramer’s V are reported to show that effect sizes are low to moderate (< 0.6), and speed of spread may not be a singularly sufficient metric for clinical use, however, these results provide evidence that the pattern of spread may be different between good and poor outcome epilepsy patients.

### E. Structural Connectivity Between Temporal Lobes is Associated with the Speed of Spread Between Regions

We hypothesized that the speed of spread between temporal lobes is associated with the structural connectivity between these lobes ^20^ – greater connectivity between temporal lobes would entail a quicker spread. Of the 71 patients, 22 acquired High Angular Resolution Diffusion Imaging (HARDI). We separated these patients with bilateral temporal lobe sampling (n = 15) and unilateral implantation (n = 7). We totaled the strength of structural connectivity between all temporal lobe regions of the left and right hemisphere (Fig 5a).

**Fig. 5.**
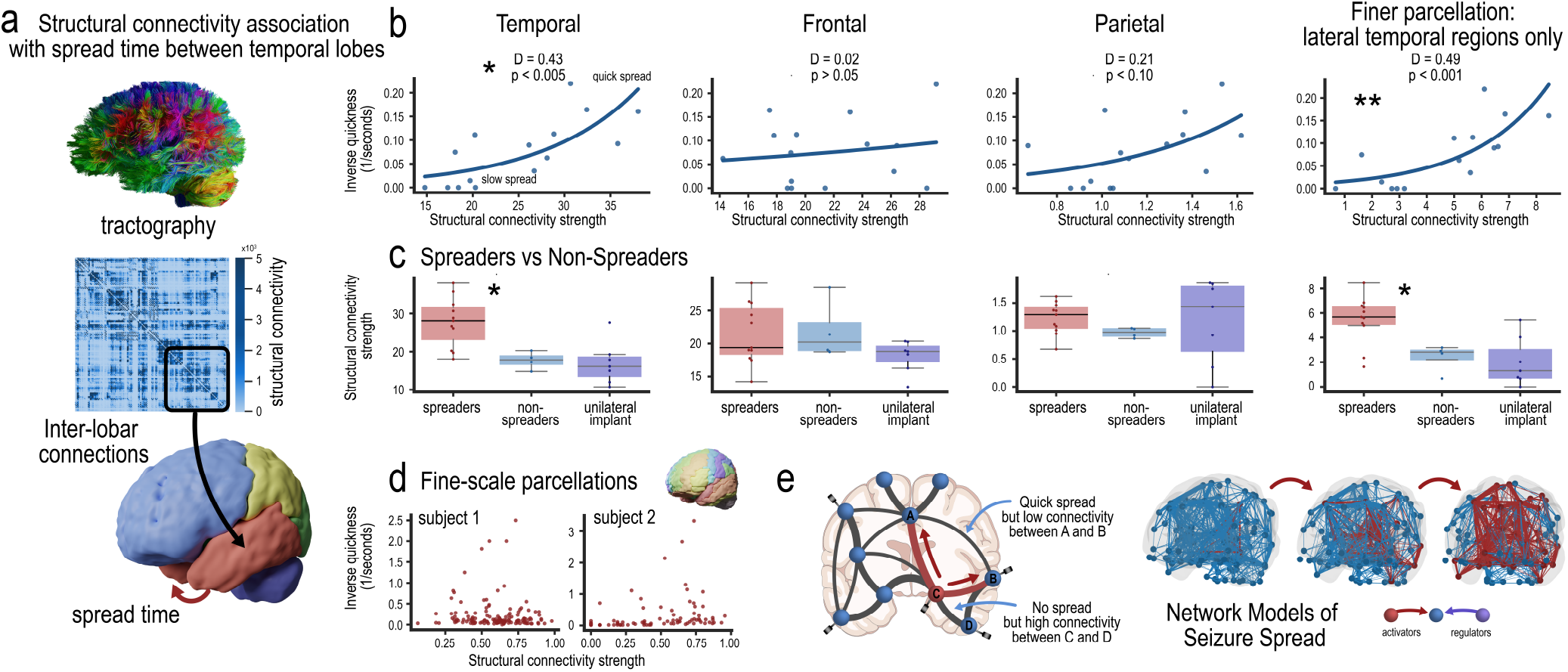
Structural Connectivity Between Temporal Lobes is Associated with the Speed of Spread Between Regions. **a**, Schematic showing how structural connectivity was measured in a subset of patients (n = 22) with High Angular Resolution Diffusion Imaging (HARDI). The heatmap represents the structural connectivity matrix of an example patient and color bar represents the strength of structural connectivity between all regions. The streamline counts between the regions of left and right lobes of each hemisphere was summed (i.e. the total streamline counts was computed between all the regions in the left and right temporal lobes). **b**, Scatter plots and a generalized linear model showing the relationship between the strength of connectivity between lobes and the timing of seizure spread between temporal lobes (n = 15 patients with bilateral sampling). X-axis indicates the total normalized streamline counts between the respective left and right lobes of each patient. Y-axis indicates inverse spread times (1/seconds). Lower numbers indicates slower spread and higher numbers indicates quicker spread. An inverse spread time of zero indicates no spread was observed between the temporal lobes of the bilaterally sampled patient. (* p < 0.005, ** P < 0.001, FDR correction with 4 tests, null hypothesis: no association between structural connectivity and spread time). **c**, Boxplots showing the structural connectivity strengths of patients (n = 22) divided into three cohorts: bilaterally sampled patients with any spread between temporal lobe structures (spreaders), bilaterally sampled patients with no spread (non spreaders), and patients who had only unilateral sampling (unilateral implant, this cohort was excluded in panel b). Mann Whitney U test was performed between the spreaders (n = 11) and non-spreader groups (n = 4). * p < 0.01 (FDR correction with 4 tests. null hypothesis: no differences in structural connectivity between spreaders and non-spreaders). Patients with unilateral implantation were plotted ad hoc and initially excluded, but not tested. **d**, An analysis of structural connectivity using a finer parcellation scale (i.e. regional versus lobar connections) of all pairwise regions. No association was found at the patient level (scatter plot of two patients) shown. **e**, Schematic showing why spread time between all pairwise regions at a finer parcellation scale may be better predicted with network models. Many regions with no spread can have high connectivity strengths. Other regions can have quick spread but low connectivity strengths.

We found a relationship in the strength of connectivity between temporal lobes and the timing of seizure spread between temporal lobes using a generalized linear model (Fig 5b, p < 0.005, n = 15 patients, percent deviance explained, *D*^2^ = 0.43, FDR corrected for 4 tests, null hypothesis: there is no relationship between the strength of structural connectivity and the speed of spread). *D*^2^ indicates the percentage of deviance explained, a generalization of the coefficient of determination *R*^2^. Here, higher connectivity strength between temporal lobes is associated with quicker spread between the temporal lobes.

We performed a negative control by computing the relationship in the spread time between temporal lobes and the strength of connectivity between other lobes. In other words, we would expect that spread time between temporal lobes is associated with the strength of structural connectivity between the temporal lobes, and spread time between temporal lobes is not determined by the strength of structural connectivity between other lobes (e.g. left frontal lobe to right frontal lobe). We found no relationship between the strength of connectivity between other lobes – frontal and parietal – and the timing of seizure spread between temporal lobes (p > 0.05, n = 15 patients, percent deviance explained, *D*^2^ = 0.02 and 0.21 for the frontal and parietal lobes respectively, FDR corrected for 4 tests, null hypothesis: there is no relationship between the strength of structural connectivity and the speed of spread).

Next we wanted to look at the strength of connectivity between smaller parcellations to explain spread time between temporal lobes. We opted to quantify the strength of structural connectivity between the lateral temporal regions (superior, middle, and inferior temporal gyri) because these regions had bilateral sampling across the 15 patients (as opposed to the hippocampus where many patients did not have bilaterally symmetric surgical placement). Additionally, smaller temporal lobe parcellations with bilateral sampling across the 15 patients preserve power. We found a relationship between the strength of connectivity between the lateral temporal lobe gyri and the timing of seizure spread between temporal lobes (p < 0.001, n = 15 patients, percent deviance explained, *D*^2^ = 0.49, FDR corrected for 4 tests, null hypothesis: there is no relationship between the strength of structural connectivity and the speed of spread). Thus, the relationship in the strength of connectivity and the timing of seizure spread between temporal lobes still holds at smaller parcellation sizes.

### F. Patients with Spread Between Temporal Lobes Have Higher Structural Connectivity than Patients With No Spread

We divided the 22 patients with structural connectivity into three cohorts: patients with any spread between the bilaterally sampled temporal lobes (n = 11), patients with no spread (n = 4), and patients with unilateral sampling who were excluded from the previous section’s analysis (n = 7). Patients who had unilateral implantation already had sufficient clinical suspicion that seizure semiology was unilateral and may not have spread to the contralateral hemisphere – they can be considered similar to the cohort with no spread.

We tested the hypothesis that patients with no spread between temporal lobes have lower structural connectivity between temporal lobes than patients with spread (Fig 5c, left box plot, Mann Whitney U test, p < 0.01 with FDR correction for 4 tests. Null hypothesis: no differences in structural connectivity between spreaders and non-spreaders). Similar to the previous section’s analysis, we found no difference in the structural connectivity between the frontal and parietal lobes in spreaders vs. non-spreader (p > 0.05, middle two box plots), and found a difference at a smaller parcellation scale by only considering the structural connectivity between the lateral temporal gyri (p < 0.01, right box plot). Patients with unilateral implantation were plotted for comparison and show similar trends in structural connectivity to patients with no spread.

### G. Structural Connectivity At Smaller Scales Cannot Predict Spread Time Between All Regions

Previous analyses focused on spread time between temporal lobes because its association with patient outcomes. We hypothesize that spread time between all pairwise regions may be predicted by the strength of structural connectivity between the pairwise regions. We found evidence that this may not be the case, and spread time instead may be associated with structural connectivity only at the lobar level (e.g. between temporal lobes), only between specific small-scale temporal lobe regions (e.g. Fig 5b and Fig 5c right most graphs), or only between unimodal association cortical areas.

We performed an analysis of structural connectivity between all pairwise regions using a finer parcellation scale (i.e. regional versus lobar connections). No association was found at the patient level (Fig 5d, scatter plot of two patients shown). We hypothesize that at smaller parcellation scales and across all pairwise regions, time of activation between pairwise regions cannot be predicted by the strength of connectivity between these regions because spread may be better predicted by network models. For example, Fig 5e shows that many regions with no spread can have high structural connectivity. Other regions can have quick spread but low connectivity strengths because spread may come from a third node activating both regions. Spread time between meso-scale regions may be better predicted by the interaction between seizure generating regions and regulatory/inhibitory regions from models that incorporate these interactions such as diffusion models, source sink models ^28^, push pull network models ^29^, Epileptor ^30^, and others.

### H. Clusters of Seizure Spread Patterns

We performed hierarchical clustering of 275 seizures across 71 patients (Fig 6). For each seizure, the pattern of seizure spread is quantified by recording the time, as a percent of seizure length, each brain region becomes active. A scatter plot of the first two principle components of seizure spread pattern is shown (Fig 6a), and each point represents a single seizure colored by cluster from the hierarchical clustering algorithm using “complete” linkage, also known as the Farthest Point Algorithm or Voorhees Algorithm ^31^.

**Fig. 6.**
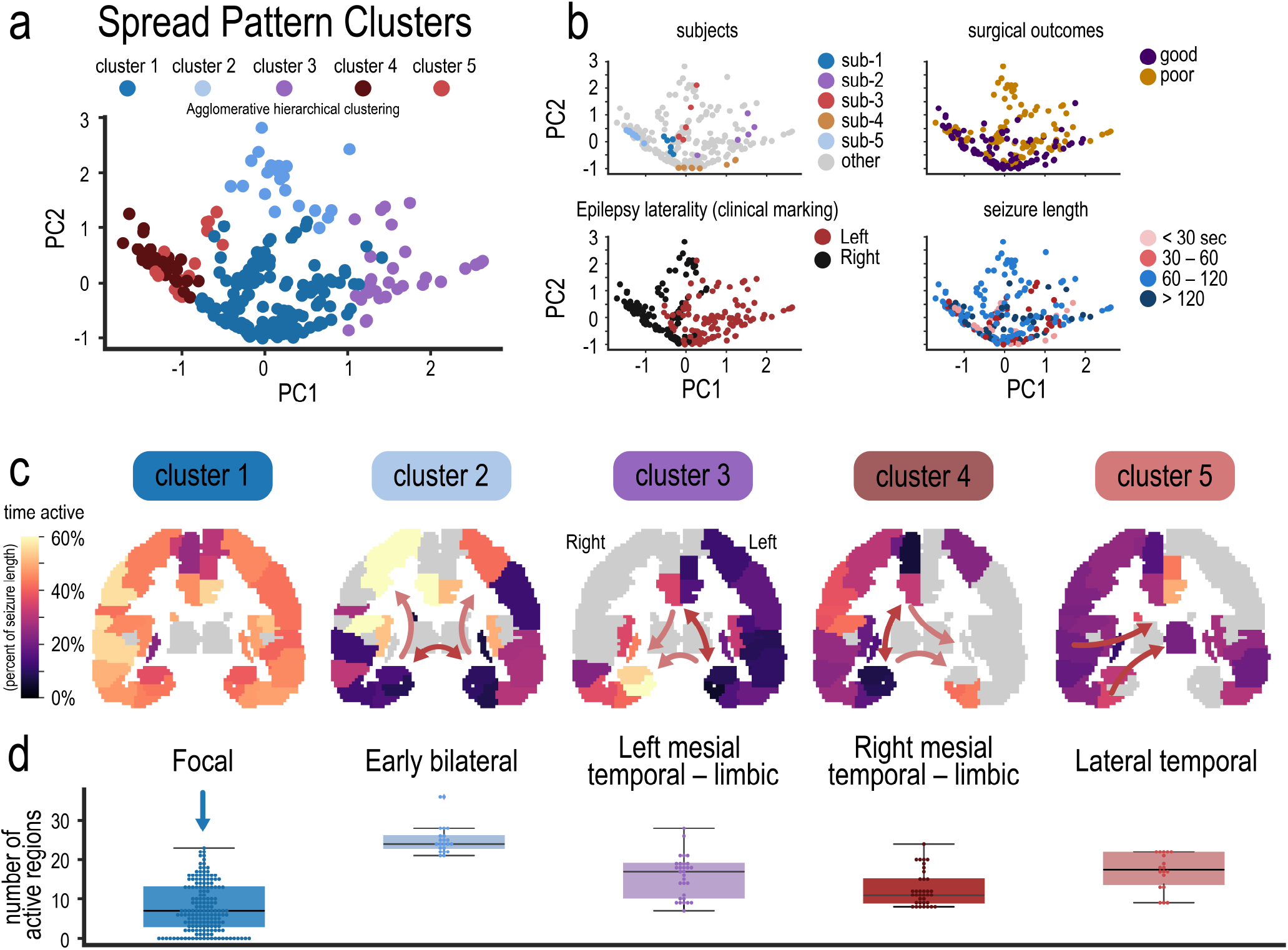
Clusters of seizure spread patterns. **a**, Hierarchical clustering of 275 seizures was performed on the pattern (location and timing) of seizure activity across 71 patients. Scatter plot of the first two principle components of seizure spread pattern is shown, and each point represents a single seizure colored by cluster. **b**, Four scatter plots are shown highlighting different attributes of the patients. Top left: five subjects with multiple seizures are highlighted. Subjects 2,3, and 4 have seizures predominantly in one cluster, but have seizures spanning multiple clusters. Top right: Seizures are highlighted by good and poor surgical outcomes. Bottom left: Seizures are highlighted by laterality of seizure onset determined though clinical chart review. Bottom right: Seizures are highlighted by length. **c**, Seizure pattern (location and timing) for each cluster is shown through a coronal slice of thee brain. Colors indicate the percentage of time in a seizure when a region was active. Darker regions indicate earlier activation. Only regions with at least two patients with seizures showing activity are colored, else regions are gray. Arrows indicate potential direction of spread. Clusters are named by the pattern of spread observed. Cluster 1 (Focal) had no early activation time, cluster 2 (early bilateral) had early activation of bilateral medial temporal lobe structures. Cluster 3 (left medial - limbic) had early activation of left medial and limbic structures. Cluster 4 (right mesial temporal - limbic) had early activation of right medial and limbic structures. Cluster 5 (lateral temporal) had early activation of right lateral temporal lobe structures. **d**, The number of regions activated at any point during a seizure for each cluster.

Four additional scatter plots show different attributes of the seizure clusters in Fig 6b. Five example patients and their seizures are highlighted. Subjects 2, 3, and 4 have seizures predominantly in one cluster, but their seizures span multiple clusters. We found patients that have seizures spanning multiple clusters usually switch between cluster 1 (focal cluster) and other clusters rather than switch between the other clusters (e.g. switch between clusters 2 and 3). Seizures are also highlighted by patients who have good or poor outcome scores at two years. Cluster 1 predominately overlaps with good outcomes. Seizures are highlighted by laterality of clinically annotated seizure onset with the principle component 1 (PC1) axis separating left and right (PC1 may separate left vs right and PC2 may separate focality or extent of spread). Finally, seizures are highlighted by the length of each seizure, with seizures < 30 seconds predominantly falling in cluster 1.

We propose a naming of each cluster based on the the timing of activity of each region averaged across the seizures in each patient and averaged across all patients within a cluster (Fig 6c). Cluster 1 (Focal) has no early activation time, cluster 2 (early bilateral) has early activation of bilateral mesial temporal lobe structures. Cluster 3 (left mesial - limbic) has early activation of left mesial and limbic structures. Cluster 4 (right mesial temporal - limbic) has early activation of right mesial and limbic structures. Cluster 5 (lateral temporal) has early activation of right lateral temporal lobe structures.

We hypothesize that Cluster 1 represents a focal, or localized, spread pattern because the average of all patients within that cluster does not indicate an early activation time of any one region. We plot the number of active regions at any time in a seizure across all the seizures in each cluster (Fig 6d). We find a lower number of active regions in the focal cluster than the other clusters. This indicates that, although seizures spread, the spread is more constrained to a lower number of regions than other clusters. A limitation is that this cluster includes patients with unilateral sampling, and thus this cluster could also include seizures in which there is not sufficient information to classify the seizure into another cluster. We further elaborate on this limitation in the “Discussion section” and why we believe the taxonomy presented in the next section is still a clinically useful representation of seizures.

### I. Taxonomy of Seizure Spread Patterns Shows the Relationship Between Clusters of Seizures

The taxonomy of seizure spread patterns using the hierarchical clustering algorithm across the 275 seizures is shown in (Fig 7). We first determined the optimal number of clusters through a principle components analysis (PCA) by plotting explained variance ratio as a function of number of components (Fig 7a). A vertical dash at n = 5 components shows a potential optimal number of components (the “elbow” method). At more clusters, we find that some clusters may be comprised of seizures from just one patient. K-means clustering is also used and sum of squared errors (SSE) is plotted as a function of number of k clusters (Fig 7b). The taxonomy of seizure spread patterns is shown in Fig 7c. Earlier branch points (e.g. the left mesial temporal branch) indicates more distinct clusters or a larger separation between clusters of other branches. The cluster numbers in the dendrogram is the same shown in Fig 6. A discussion and interpretation of these clusters and branch points are in the “Discussion section”.

**Fig. 7.**
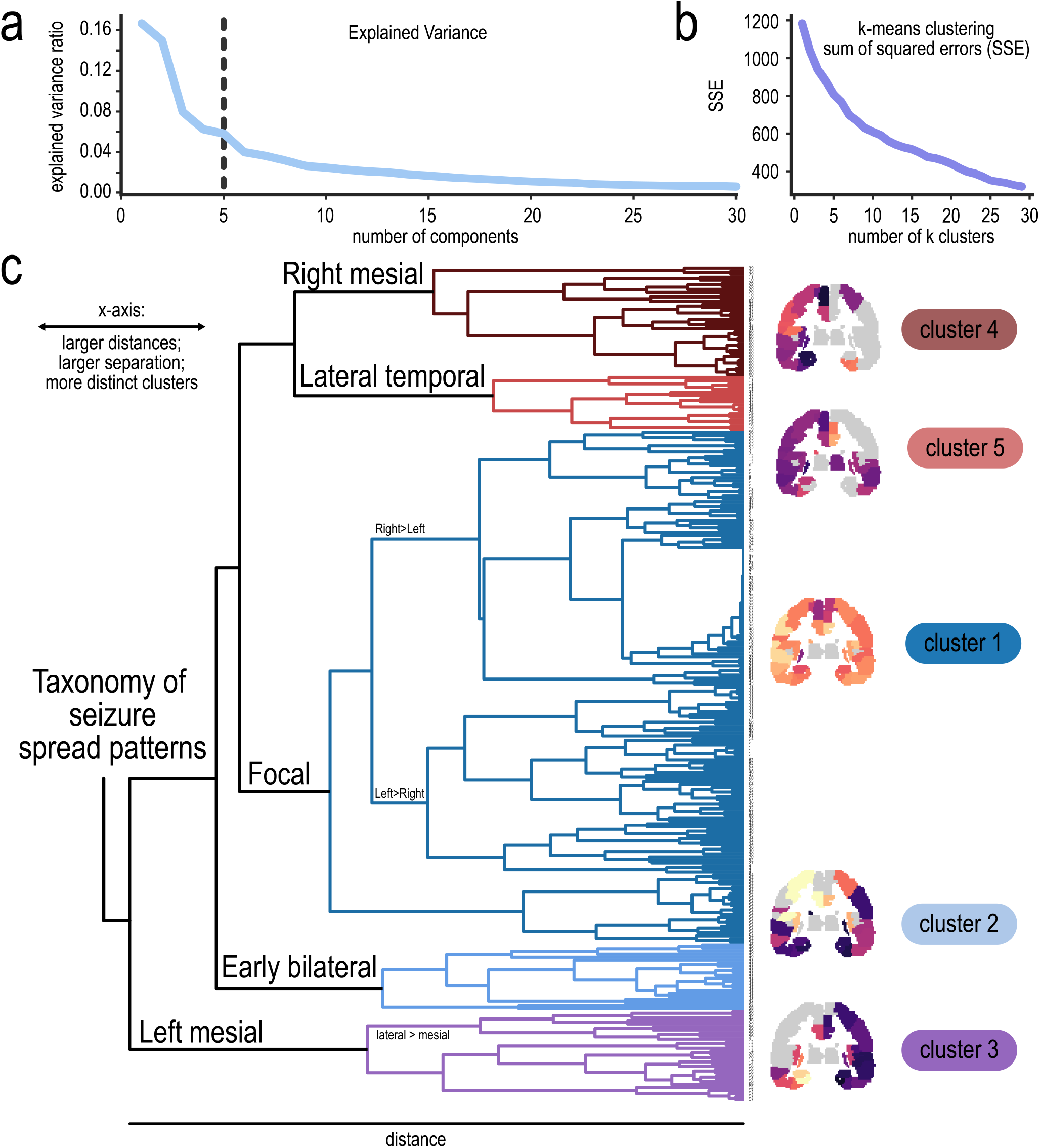
Taxonomy of Seizure Spread Patterns. **a**, Principle components analysis (PCA) showing explained variance ratio as a function of number of components to select for an optimal number of clusters. A vertical dash is at n = 5 components **b**, K-means clustering sum of squared errors (SSE) is plotted as a function of number of k clusters. **c**, The taxonomy of seizure spread patterns is shown using hierarchical/agglomerative clustering. Colors of branches correspond to clusters from Fig. 6. X-axis shows the euclidean distance between clusters with a “complete” linkage function, also known as the Farthest Point Algorithm or Voorhees ^31^ Algorithm. For example, the left mesial cluster is the first branch point indicating that this spread pattern is a distinct evolution of seizure spread and is most different from the other clusters.

## Discussion

In this study, we develop, validate, and compare different algorithms to measure seizure spread with the goal to organize and classify hierarchical patterns of seizure spread. We find that deep learning algorithms are highly effective in differentiating ictal and interictal states over single features (Fig 2) which can be used to detect seizure onset and measure spread (Fig 3). We discover that poor outcome patients have more distributed regions involved in seizures (Fig 4a-d) and seizure spread within 5 seconds between the average activation time of the left and right temporal lobes yields a specificity of 94% in differentiating good and poor outcome surgical patients at two years (42% sensitivity, Fig 4e). This speed of seizure spread between temporal lobe regions is associated with the strength of structural connectivity between temporal lobes, but not between other regions (Fig 5). Finally, hierarchical clustering over 275 seizures and 71 patients shows 5 distinct clusters and the relationship between these clusters (Fig 6 and (Fig 7)). We name each of the 5 major clusters – Cluster 1: focal, Cluster 2: early bilateral, Cluster 3: left mesial temporal - limbic, Cluster 4: right mesial temporal - limbic, and Cluster 5: lateral temporal.

### A. The Focus of Seizure Activity Past Seizure Onset

In our study, we show the pattern of seizure activity may be an important marker that can predict response to epilepsy surgery. While correct identification of seizure onset contacts is essential for the success of surgery and has been a major focus in computational studies attempting to identify location of seizure onset, it perhaps should not be the only focus of EEG interpretation nor the focus to identify ideal surgical candidates and brain regions targeted for surgery. Here, we show that the pattern of spread, whether through the extent of spread (Fig 4a-d) or speed of spread (Fig 4e) is associated with surgical outcomes at two years.

Furthermore, the ability to quantify patterns of seizure spread – whether through complicated deep learning algorithms or though simple features – opens new avenues to study epilepsy patholophysiology and seizure evolution. Although we present evidence that deep learning algorithms may be superior in capturing spread patterns over simple features such as line length, these singular features still capture onset and spread better than chance (Fig 3), and many labs or clinical software may be suited for reporting spread patterns. For example, the patient in Fig 4a shows 14 seizures captured during their hospital stay and we can observe how the pattern of spread changes during the days to weeks a patient may stay in the epilepsy monitoring unit. We can quantify whether a seizure is a stereotypical pattern and is representative of their semiology to be used for interpretation and localization of seizure onset. We can also observe how certain external factors, such as medications, sleep deprivation, and other habits may change the patterns of seizure activity over time. We found that these seizure spread algorithms work on both ECoG and SEEG implantations, so they may also have utility in scalp EEG, which benefits from standard sampling across patients (although more coarse in localization than intracranial implantations).

### B. The Hierarchical Organization of Seizure Spread Patterns

The goal of hierarchical clustering is to find the overarching organization and classification of a data set in an unsupervised manner ^32^. Here, we organize the patterns of seizure spread into distinct clusters, and the discussion here is to provide interpretation of the unsupervised learning algorithm. The dendrogram of Fig 7 shows the relationship between the clusters of seizure spread patterns. Earlier branch points indicate a large separation from other clusters.

The left mesial temporal cluster (cluster 2) is the first branch point indicating that this spread pattern is a distinct evolution of seizure spread and is most different from the other clusters. We interpret this as seizure activity with left mesial temporal involvement and spread is a distinct form of seizure pathophysiology and evolution. Other seizures with left mesial temporal seizures are included in cluster 1 (the focal cluster), however hierarchical clustering indicates that this subset of left mesial temporal involvement is more limited, and that left mesial temporal involvement *with* spread (cluster 3) may be a distinct form of seizure spread pattern. Some patients with left mesial temporal onset switched between clusters 1 and 3, and we interpret this switching as evidence that the exact etiology of each seizure in a patient may not necessarily be the same. Epilepsy pathophysiology may change from seizure to seizure (e.g. regulatory mechanisms, excitatory/inhibitory responses, push pull networks, etc. may change across time during a patient’s hospital stay), and this may provide clues to a clinician how to plan treatment for their patient.

At n = 5 clusters, we did not see a distinct separation of left mesial and lateral temporal lobe clusters as in clusters 4 and 5 (right sided mesial temporal and lateral temporal patterns, respectively). However, investigation into some of the branches in cluster 3 did show a branch with earlier left lateral temporal lobe activation than left mesial temporal lobe activation (lateral > mesial). To see this separation would require an increase in clusters *a priori*, but more than 5 clusters would result result in some clusters (particularly in cluster 1) being comprised of only one patient. Thus separation of the left mesial branch into more distinct clusters could not be done systematically.

Overall, we find that this hierarchical clustering separation also aligns similarly to clinical investigation of epilepsy onset – is it left or right sided, does it have early bilateral activation or more focal spread with limited regions, is is lateral or mesial temporal lobe epilepsy?

### C. Limitations

A major limitation to this study is that the organization found in the hierarchical clustering can be affected by the implantation and sampling bias of our patients ^33^. For example, the occipital lobe is rarely implanted, and thus clustering using this region provides little discriminating information to the clustering algorithms. Another example is that through principle components analysis (Fig 6a), PC1 seems to differentiate spread patters by laterality – spread from the right hemisphere has negative PC1 value, spread from the left has positive PC1 values, and bilateral spread (namely cluster 2) has PC1 values close to zero. We did not see a principle component that organizes spread patters in an anterior-posterior brain axis probably because the focus of implantation is heavily subject to a left-right organization.

Furthermore, cluster 1 is the focal or localized cluster, and patients with unilateral sampling fall into this cluster perhaps because seizures captured in these patients may not have sufficient information to classify the seizure into another cluster (i.e. the spread pattern is classified into the focal cluster because there is little information about seizure spread to other brain region in a unilateral implantation).

Despite this limitation, however, we find that many patients with unilateral sampling or other implantation schemes (rather, the lack of certain sampling from regions like the occipital lobe) have their respective implantation schemes *for a clinical reason* – there is evidence that seizure activity may be limited to the regions targeted for implantation *a priori*. The patterns of spread classified by physicians are typically implanted in a stereotyped fashion (i.e. patients with suspected left mesial temporal lobe epilepsy largely have similar structures targeted for implantation with modifications based on clinical history and other findings). For example, many patients with limited sampling and spread (Fig 6b) have good outcomes not because their sampling was limited *per se*, but rather because their spread pattern was predicted to be well-localized by their physicians, they were implanted to confirm seizure onset, and subsequently had a good outcome after surgery because seizure onset was already biased in its focal localization.

Thus we believe that in this study, our patients’ implantation and the activity recorded in those regions are a fair representation of the spread pattern within the brain. Patients across institutions and other studies may have similar sampling and implantation schemes to our cohort of 71 patients and the hierarchical organization found in this study may provide a fair representation of the taxonomic organization of seizure spread patterns in a clinically relevant manner.

## Conclusion

The pattern of seizure activity past seizure onset may help direct treatment of refractory epilepsy patients and can indicate if surgical intervention or other treatment options may have the best chance to improve a patient’s quality of life. We propose a shift in epilepsy research from a primary focus in identification of seizure onset to quantifying the patterns of seizure activity past onset.

## Materials and Methods

### A. Clinical Data and outcome scoring

Seventy-one individuals (mean age 33 ± 12; 31 female) underwent intracranial EEG implantation (iEEG) of either electrocorticography (ECoG, n = 23) or stereoelectroencephalography (SEEG, n = 48, Supplementary Table S1). Across the 71 patients, 275 seizures were captured (Fig S1, mean length 85 ± 94 seconds; mean number of seizures captured per patient 3.9 ± 3.8 seizures). Fifty eight patients had Engel outcome scores at two years after undergoing epilepsy surgery. Engel I outcome scores were classified as good outcomes and Engel II-IV were classified as poor outcomes.

### B. Intracranial EEG Acquisition

ECoG and SEEG electrodes were implanted in patients based on clinical necessity. Continuous intracranial EEG (iEEEG) signals were obtained for the duration of each patient’s stay in the epilepsy monitoring unit. Intracranial data was recorded at 256, 512, or 1024 Hz for each patient. Seizure onset times were defined by the unequivocal electrographic onset (UEO) ^34^. Interictal data were taken at least six hours before seizure onset and were 180 seconds in length. All annotations were verified by neurologists and consistent with detailed clinical documentation. The spacing between SEEG contacts is 5 mm and the contacts are 2.41 mm in size.

### C. Electrode Localization

In-house software ^35^ was used to assist in localizing electrodes after registration of pre-implant and post-implant images (T1w and CT images). All electrode coordinates and labels were saved and matched with the electrode names on IEEG.org. All electrode localizations were verified by a board-certified neuroradiologist (J.S.).

### D. Pre-processing of EEG

Following removal of artifact-ridden electrodes, iEEG signals were bipolar referenced. Signals were notch-filtered at 60 Hz to remove power line noise and low-pass and high-pass filtered at 127 Hz and 1Hz to account for noise and drift. iEEG signals were downsampled to 128 Hz because a larger sampling rate would not fit into memory of a GPU for training and testing of deep learning algorithms. Signals were then pre-whitened using a first-order autoregressive model to account for slow dynamics. All iEEG signals for each channel were normalized to each respective channel’s interictal data. This was done by applying the Python package sklearn robust scaler function. This function scales features by removing the median and scales the data according to the interquartile range.

### E. Deep learning algorithms

The structure of the deep learning algorithms and their parameters as as follows. Python packages and versions are listed at the end of the Methods section. Python code can be found at https://github.com/andrewyrevell/revellLab/ in the SeizureSpread package. The code for the deep learning algorithms is provided explicitly below because these algorithms are central to measuring seizure spread.

#### E.1. Global parameters

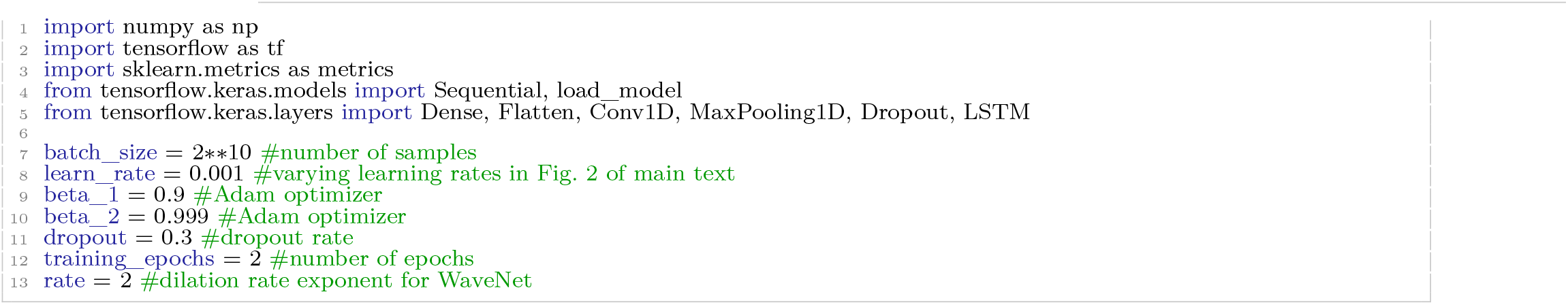

#### E.2. WaveNet

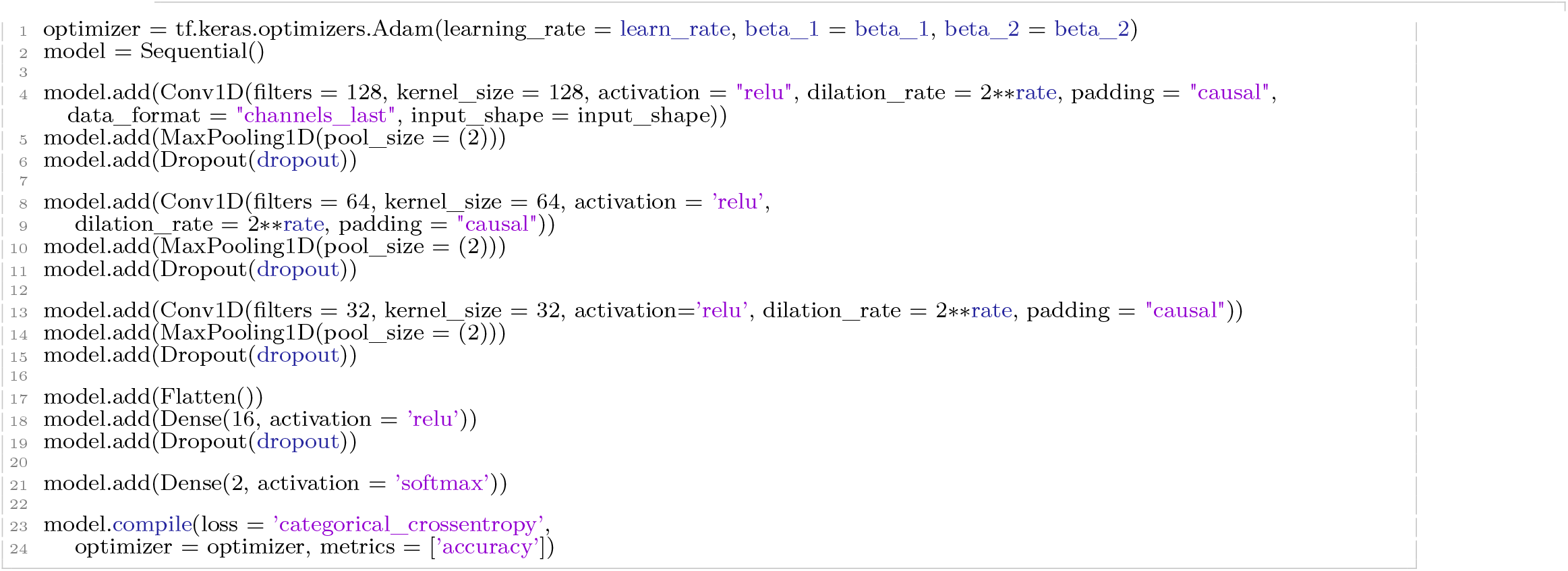

#### E.3. 1D CNN

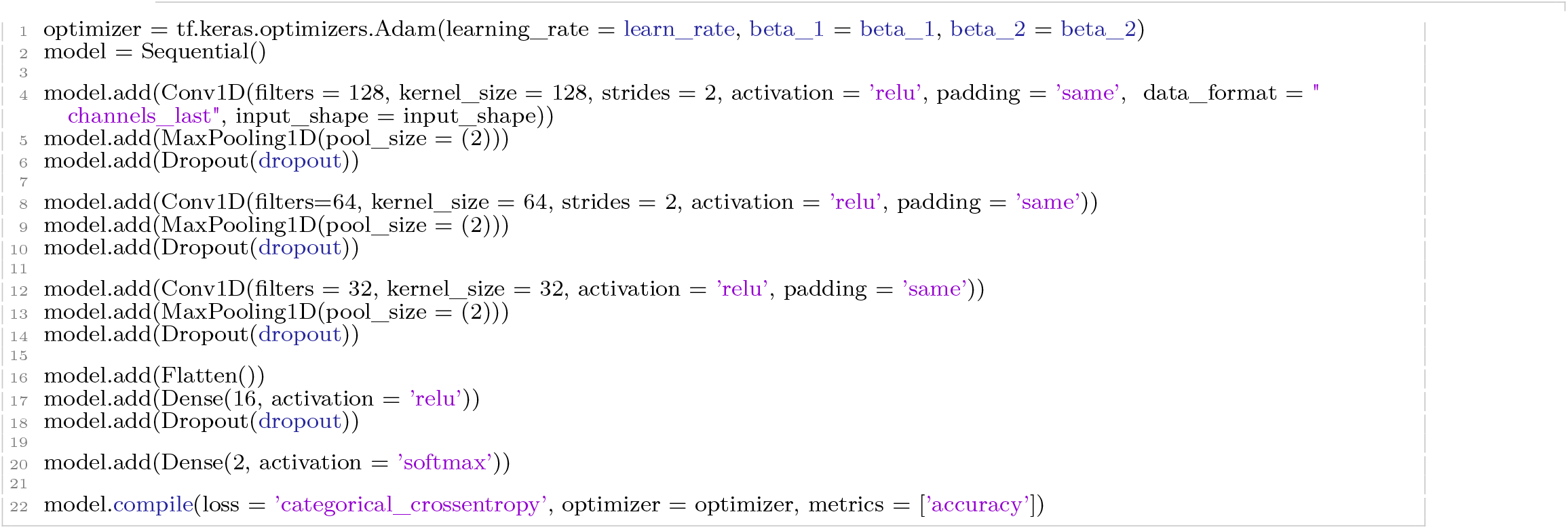

#### E.4. LSTM

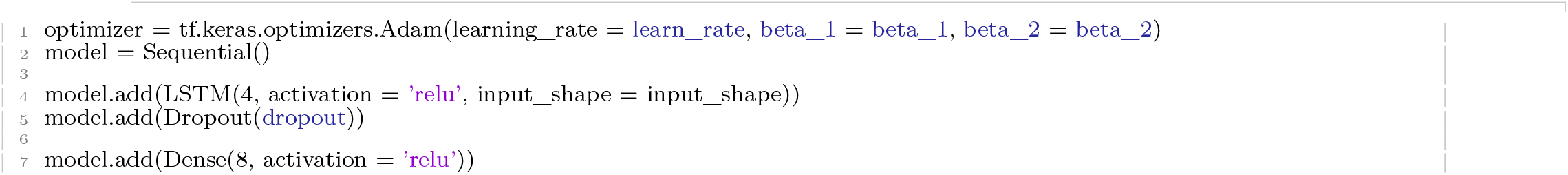
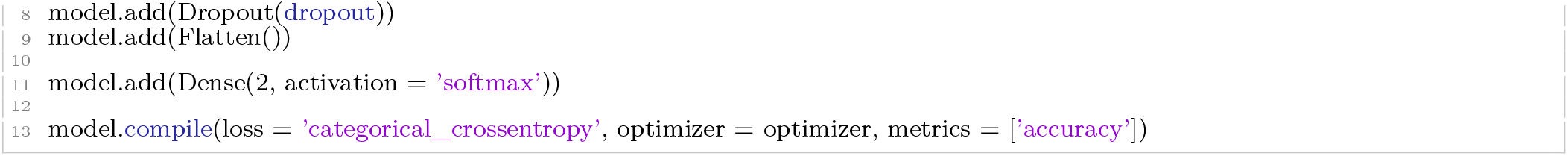

### F. Single features - Absolute Slope, Line Length, Broadband Power

The single features, absolute slope ^21^, line length ^22,23^, and broadband power were calculated on the pre-proceessed EEG data. Broadband power was calculated using the Scipy Python package version 1.5^36^, and the function scipy.signal.welch (default parameters, with FFT epoch length equal to 1s).

Line length:

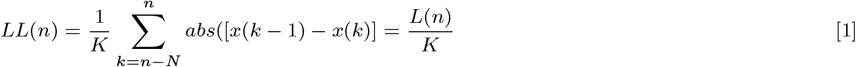

where LL(n) is the normalized line length value at a discrete time index n. L(n) is the sum of distances between successive points within the sliding window of size N sample points. x[k] is the value at the kth sample. K is the normalization constant. ^22^.

Absolute slope:

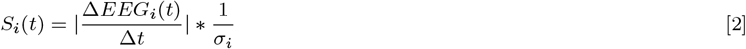

where i runs over all channels, t denotes time, and *σ_i_* denotes the standard deviation of *S_i_*(*t*) during the interictal period of channel *i*.

### G. Training and Testing, and Measuring Seizure Spread

Training and testing data were broken into 1 second windows with 0.5 seconds overlap on iterictal data (”definitely not seizing”) and sections of ictal data on each channel that an annotator determined to be “definitely seizing.” Performance was quantified with area under the curve (AUC) for differentiating interictal and “definitely seizing” windows. A leave-one-out cross validation approach was used on n = 13 patients.

To quantify seizure spread after training and testing, 180 seconds of preictal and 180 seconds of postictal data were collected in addition to the seizure. For each channel, 1 second windows with 0.5 seconds overlap were used to calculate probability of seizure (for the deep learning algorithms) or the normalized single feature. Probabilities or single features were smoothed over 20 seconds. Onset of activity for each channel was determined by the time window at which the smoothed value crossed a pre-determined set threshold after the unequivocal onset.

### H. Validation of Seizure Spread Algorithms: Median Rank Percent of Seizure Onset Contacts

After measuring seizure spread, the performance of each algorithm was assessed by its ability to appropriately rank seizure onset contacts with physician markings. The agreement between physician markings and each algorithm marking is calculated, and the algorithm ranking of the seizure onset contacts were averaged (median). This median ranking is divided by the total number of implanted contacts to normalize for differences in implantations. Note, this calculation penalizes ranking scores for physicians who marked large number of contacts. However, we focused on comparing algorithms, and this penalty is equal between algorithms.

### I. Atlas choice

We chose the Automated Anatomical Labeling (AAL) atlas ^37–39^ in this study because (1) it is a common structural atlases used to create structural connectivity (2) it contains regions with sufficient depth to include depth electrodes (contacts that fall outside the atlas are excluded from analysis, reducing power), and (3) the AAL atlas provides appropriate power to study the structure-function relationship of the brain ^40^ (i.e. its parcellation scheme is appropriate to use in studies incorporating both structural data from diffusion imaging and functional data from iEEG. Adding additional atlases may reduce power of our study).

### J. Calculating Extent of Spread

The extent of seizure spread can be quantified in two ways: (1) by the number of contacts and (2) by the number of brain regions activated over time (Fig 4). Each metric was converted into a percentage by (1) dividing the number of active contacts by the total number of contacts implanted (excluding contacts that fell outside the brain or artifact contacts) and (2) dividing the number of active brain regions by the total number brain regions sampled. If multiple contacts fell in a brain region, then that regions was still counted only once. The activity (i.e. probabilities or single feature metrics) of all the contacts within a single region were averaged together. That region was considered active if the average probability or metric fell above a predetermined threshold.

### K. Calculating Speed of Spread

Activation times of all contacts in the temporal lobe were averaged together for each of the left and right hemispheres to calculate spread time. The absolute value difference in the average activation times was recorded in seconds. To account for seizure with no spread, inverse spread times were calculated (1/spread time). For example, spread time could not be calculated or would be considered infinite if the left temporal lobe was active and the right temporal lobe never became active. Therefore, inverse spread time would be adjusted to zero and could be compared to seizures with spread.

### L. Contingency Tables, Sensitivity, and Specificity

Cutoff times at 5, 10, 15, 20,30, and 60 seconds were used to differentiate good and poor outcomes. Outcomes at 2 years were used. Sensitivity, specificity, positive predictive values (PPV), negative predictive values (NPV), chi-square test, and Cramer’s V are reported for each cutoff.

### M. Structural Connectivity

The below subsections detail the methodology for calculating structural connectivity.

#### M.1. Imaging protocol

Prior to electrode implantation, MRI data were collected on a 3T Siemens Magnetom Trio scanner using a 32-channel phased-array head coil. High-resolution anatomical images were acquired using a magnetization prepared rapid gradient echo (MPRAGE) T1-weighted sequence (repetition time = 1810 ms, echo time = 3.51m, flip angle = 9, field of view = 240mm, resolution = 0.94×0.94×1.0 mm3). High Angular Resolution Diffusion Imaging (HARDI) was acquired with a single-shot EPI multi-shell diffusion-weighted imaging (DWI) sequence (116 diffusion sampling directions, b-values of 0, 300, 700, and 2000s/mm2, resolution = 2.5×2.5×2.5 mm3, field of view = 240mm). Following electrode implantation, spiral CT images (Siemens) were obtained clinically for the purposes of electrode localization. Both bone and tissue windows were obtained (120kV, 300mA, axial slice thickness = 1.0mm)

#### M.2. Diffusion Weighted Imaging (DWI) Preprocessing

HARDI images were subject to the preprocessing pipeline, QSIPrep, to ensure reproducibility and implementation of the best practices for processing of diffusion images ^41^. Briefly, QSIPrep performs advanced reconstruction and tractography methods in curated workflows using tools from leading software packages, including FSL, ANTs, and DSI Studio with input data specified in the Brain Imaging Data Structure (BIDS) layout.

#### M.3. Structural Network Generation

DSI-Studio (http://dsi-studio.labsolver.org, version: December 2020) was used to reconstruct the orientation density functions within each voxel using generalized q-sample imaging with a diffusion sampling length ratio of 1.25^42^. Deterministic whole-brain fiber tracking was performed using an angular threshold of 35 degrees, step size of 1mm, and quantitative anisotropy threshold based on Otsu’s threshold ^43^. Tracks with length shorter than 10mm or longer than 800mm were discarded, and a total of 1,000,000 tracts were generated per brain. Deterministic tractography was chosen based upon prior work indicating that deterministic tractography generates fewer false positive connections than probabilistic approaches, and that network-based estimations are substantially less accurate when false positives are introduced into the network compared with false negatives ^44^. To calculate structural connectivity, the AAL atlas was used. Structural networks were generated by computing the number of streamlines passing through each pair of atlas regions. Streamline counts were log-transformed and normalized to the maximum streamline count, as is common in prior studies ^45–48^. For each left and right hemisphere all the temporal lobe, frontal lobe, and parietal lobe structures were combined and the structural connectivity between each hemisphere of each lobe were totaled together to represent the structural connectivity between the hemispheres of each lobe.

### N. Generalized Linear Models to Quantify the Relationship Between Structural Connectivity and Speed of Spread

The Statsmodel Python package was used to construct a Tweedie regressor with 1.1 power. Structural connectivity was the independent variable and inverse spread time was the dependent variable. The percent deviance explained, *D*^2^, was calculated using the Sklearn.linear_model Python package and the TweedieRegressor score method. *D*^2^ indicates the percentage of deviance explained, a generalization of the coefficient of determination *R*^2^.

### O. Hierarchical Clustering Algorithm

The Scipy ^36^ python package is used to calculate hierarchical clustering: Scipy.cluster.hierarchy.linkage with method = “complete”. This method is also known as the “Farthest Point Algorithm” or Voorhees Algorithm ^31^. This algorithm was chosen because it defines the distance between two groups as the distance between the two farthest-apart members. The advantage is it usually yields clusters that are well separated and compact. The default “single” method (also known as the “nearest neighbor method” did not yield interpretable results; the majority of seizures fell in one cluster with large number of clusters contained of single seizures without a clear hierarchical organization. Clustering was performed on a matrix of shape 275 x 120, where 275 represents the number of seizures in this study and 120 represents the number of regions in the AAL atlas. Each cell in the matrix contained the percent of time into a seizure that a region became active.

### P. Python Packages and Versions

The conda environment for the analyses can be ofound in https://github.com/andyrevell/revellLab/ in the envirnoments folder. The YAML file is below:

#### P.1. Conda environment YAML file

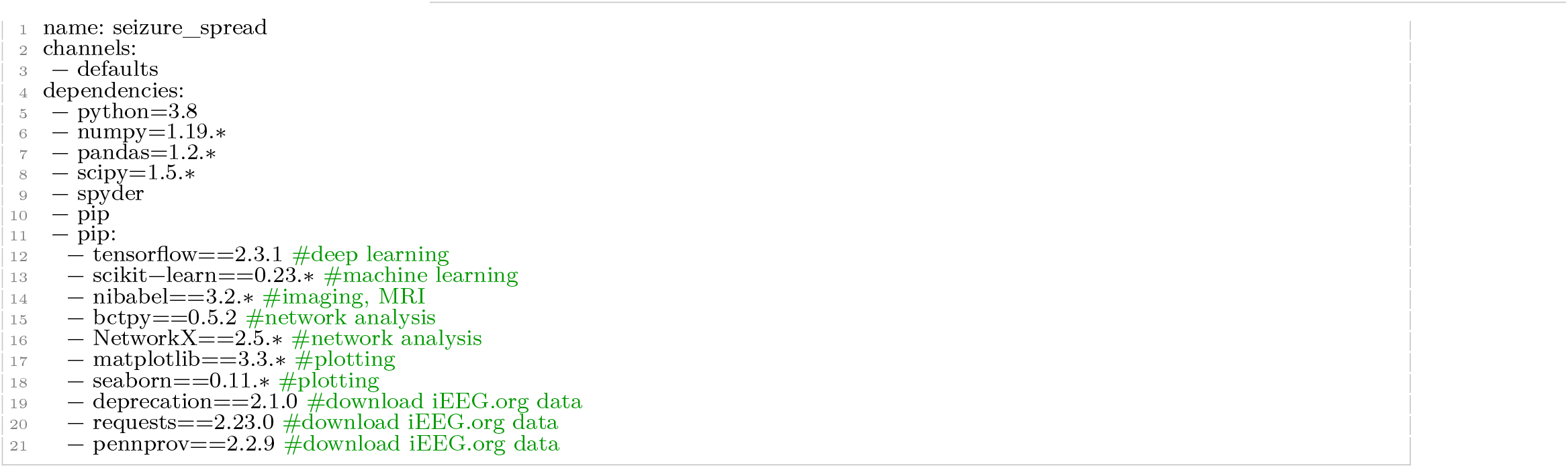

## Acknowledgements

We thank Adam Gibson, Carolyn Wilkinson, Jacqueline Boccanfuso, Magda Wernovsky, Ryan Archer, Kelly Oechsel, members of Andrew’s Thesis Committee, Braden Kelner (for editing), and all other members and staff of the Center for Neuroengineering and Therapeutics for their continued help and support in this work.

## Supplementary Materials

Please see supplemental materials below.

• Figures

– Fig. S1: Distribution of Seizure Lengths and Number Per Patient
– Fig. S2: Seizures Colored By Other Attributes
– Fig. S3: Effect Size Comparisons Between Seizure Detection Algorithms in Extent and Speed of Spread

The figures below contain supplemental information for the main text.

**Fig. S1.**
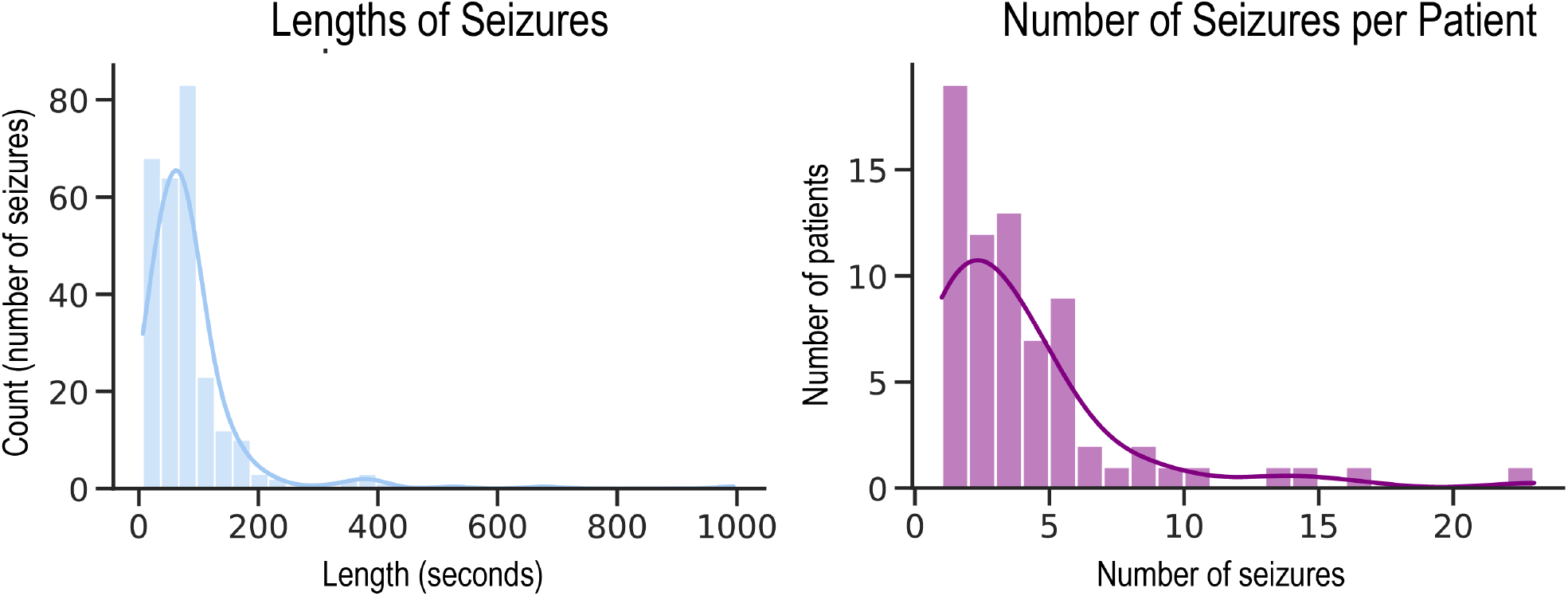
Distribution of Seizure Lengths and Number Per Patient. Left: The distribution of seizure lengths across all 275 seizures in this study. Mean: 85 seconds, median: 68 seconds, sd: 94 seconds. Right: The distribution of the number of seizures per patient. Mean: 3.9, median: 3.0, sd: 3.8.

**Fig. S2.**
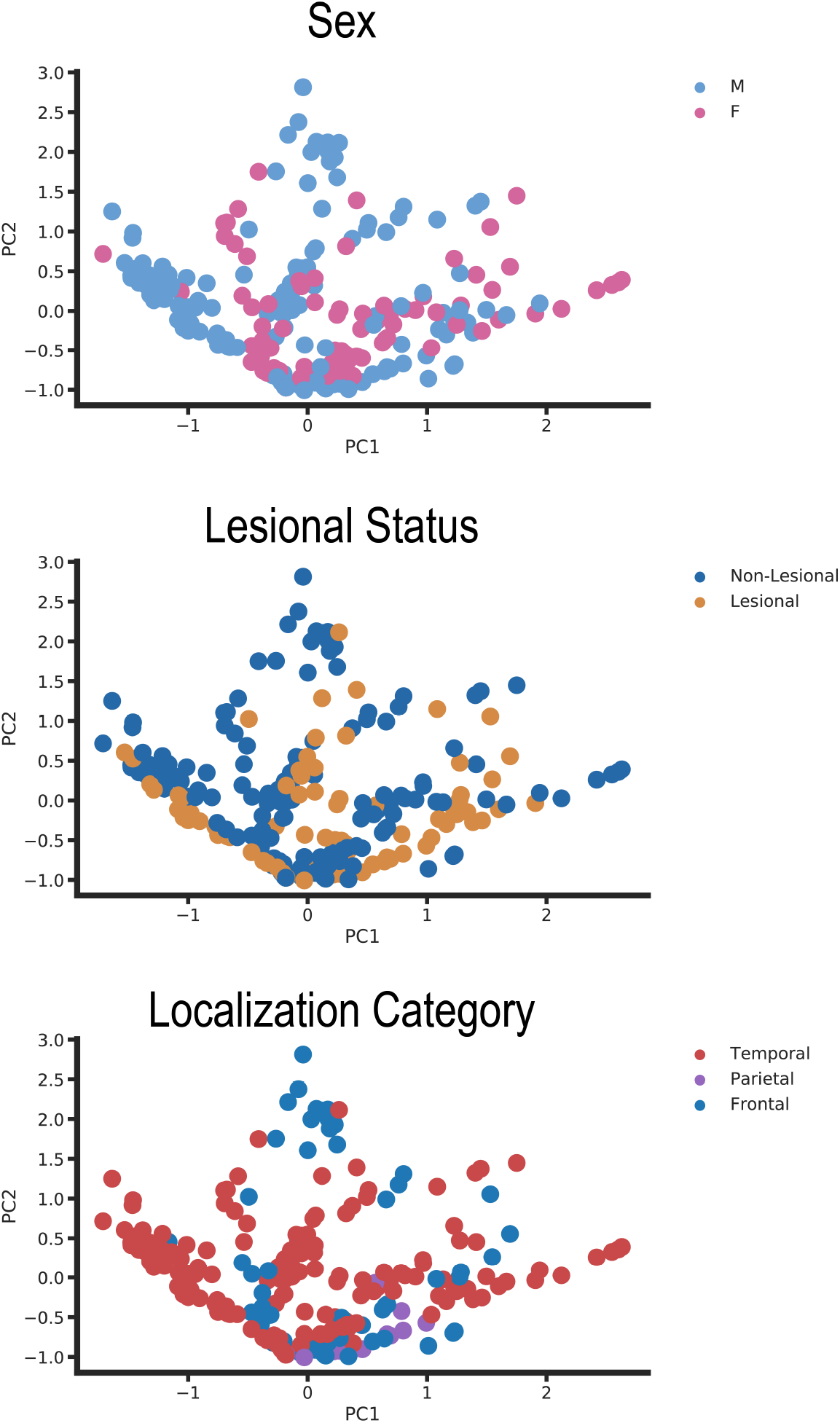
Seizures Colored By Other Attributes. Seizures from Fig. 6 are colored by other attributes such as sex (top), lesional status (middle), and lobar localization (bottom).

**Fig. S3.**
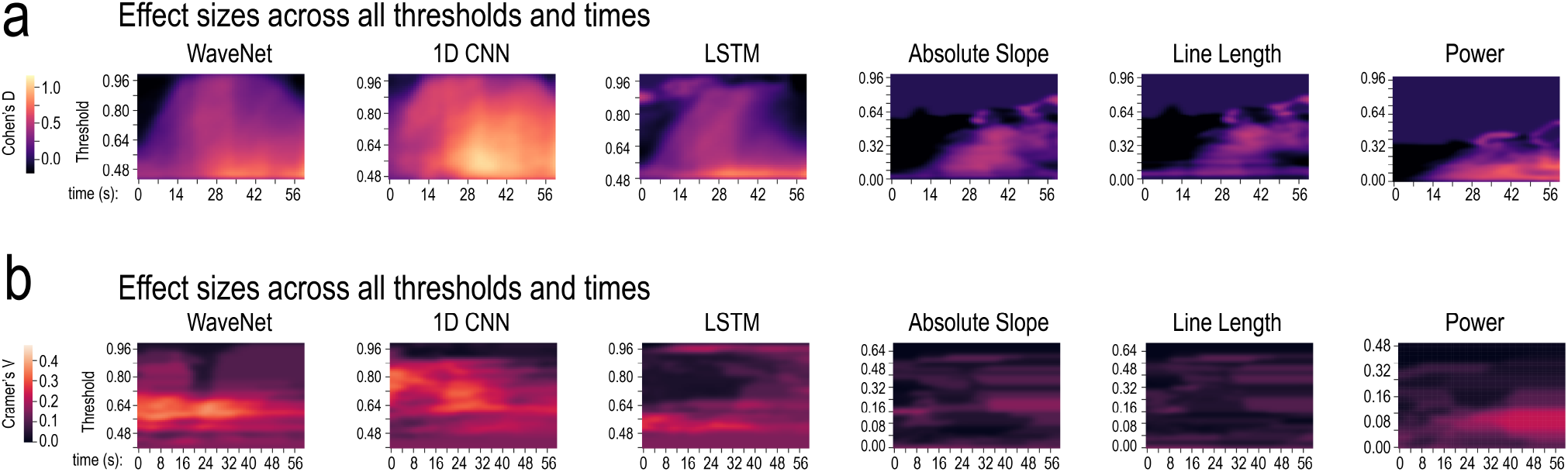
Effect Size Comparisons Between Seizure Detection Algorithms in Extent and Speed of Spread. **a**, Effect sizes across all thresholds and times for comparing the extent of seizure spread in good a poor outcome patients. Heatmaps and color bars represent Cohen’s D. **b**, Effect sizes across all thresholds and times for comparing the association between the speed of seizure spread between temporal lobe regions and surgical outcomes. extent of seizure spread in good a poor outcome patients. Heatmaps and color bars represent Cramer’s V.

