## Supplemental File for "A Taxonomy of Seizure Spread Patterns, Speed of Spread, and Associations With Structural Connectivity"

### Supplementary Materials

Please see supplemental materials below.

- Figures
  - Fig. S1: Distribution of Seizure Lengths and Number Per Patient
  - Fig. S2: Seizures Colored By Other Attributes
  - Fig. S3: Effect Size Comparisons Between Seizure Detection Algorithms in Extent and Speed of Spread

The figures below contain supplemental information for the main text.

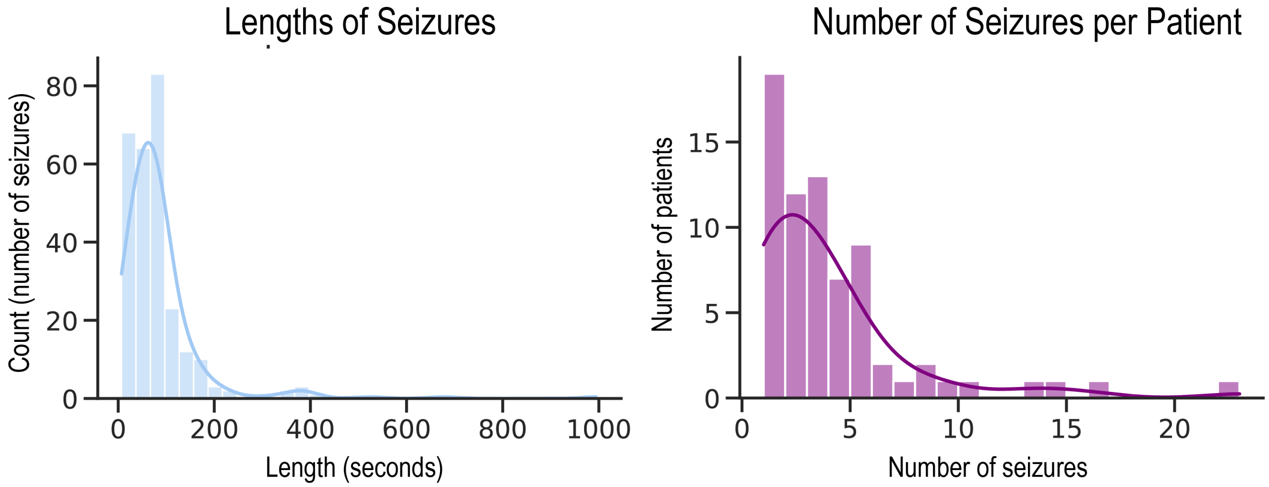


**Fig. S1. Distribution of Seizure Lengths and Number Per Patient.** | Left: The distribution of seizure lengths across all 275 seizures in this study. Mean: 85 seconds, median: 68 seconds, sd: 94 seconds. Right: The distribution of the number of seizures per patient. Mean: 3.9, median: 3.0, sd: 3.8.

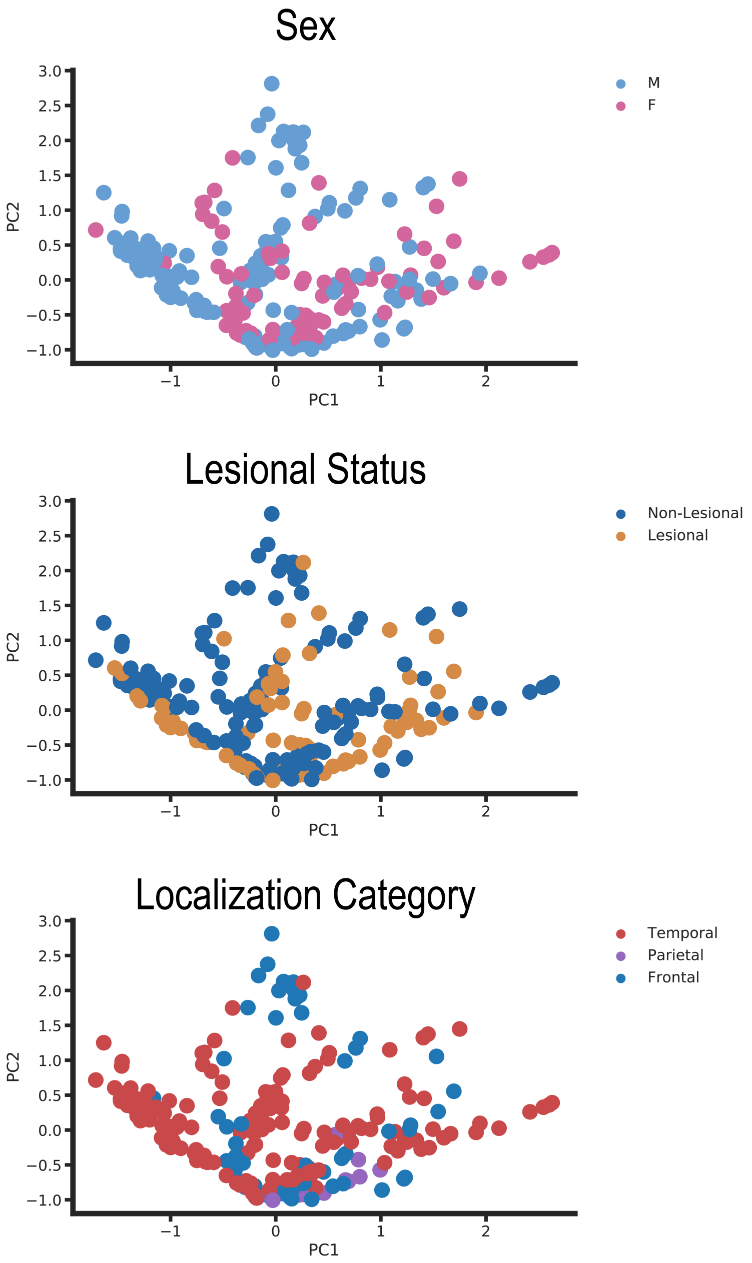


**Fig. S2. Seizures Colored By Other Attributes.** | Seizures from Fig. 6 are colored by other attributes such as sex (top), lesional status (middle), and lobar localization (bottom).

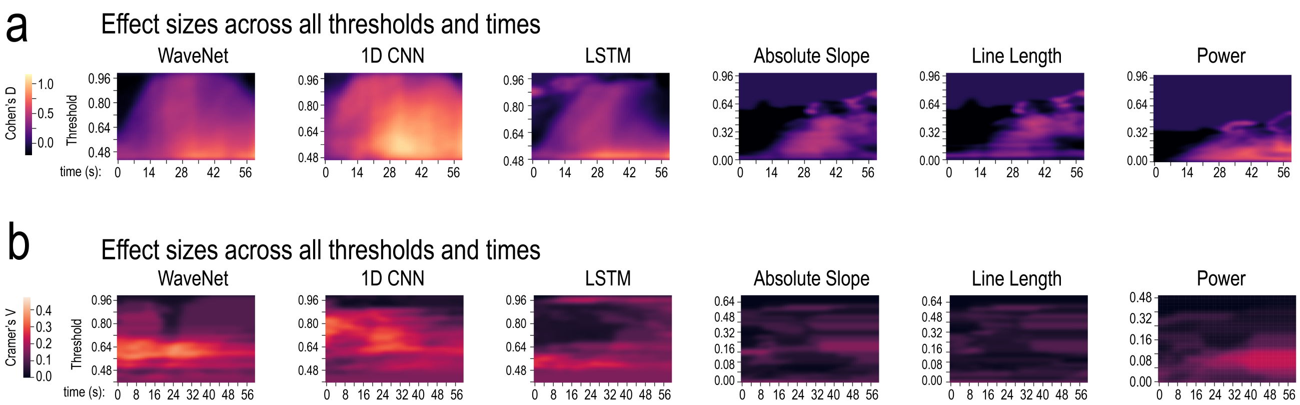


**Fig. S3. Effect Size Comparisons Between Seizure Detection Algorithms in Extent and Speed of Spread.** | **a**, Effect sizes across all thresholds and times for comparing the extent of seizure spread in good a poor outcome patients. Heatmaps and color bars represent Cohen's D. **b**, Effect sizes across all thresholds and times for comparing the association between the speed of seizure spread between temporal lobe regions and surgical outcomes. extent of seizure spread in good a poor outcome patients. Heatmaps and color bars represent Cramer's V.
